# Tracking the misallocation and reallocation of spatial attention towards auditory stimuli

**DOI:** 10.1101/2023.11.25.568671

**Authors:** Ananya Mandal, Anna M. Liesefeld, Heinrich R. Liesefeld

**Affiliations:** General and Experimental Psychology, Ludwig-Maximilians-Universität Munich, 80802, Munich, Germany; Graduate School for Systemic Neurosciences, LMU Munich, 82152, Planegg, Germany; Department of Psychology, Universität Bremen, 28359 Bremen, Germany

**Keywords:** auditory search, auditory distraction, N2ac, ERP component latency, auditory attention capture, selective attention

## Abstract

Completely ignoring a salient distractor presented concurrently with a target is difficult and sometimes attention is involuntarily attracted to the distractor’s location (*attentional capture*). Employing the N2ac component as a marker of attention allocation towards sounds, in this study we investigate the spatio-temporal dynamics of auditory attention across two experiments. Human participants (male and female) performed an auditory search task, where the target was accompanied by a distractor in 2/3rd of the trials. For a distractor more salient than target (Exp. 1), we observe not only a distractor N2ac (indicating attentional capture), but the full chain of attentional dynamics implied by the notion of attentional capture, namely: (a) the distractor captures attention before the target is attended, (b) allocation of attention to the target is delayed by distractor presence, and (c) the target is attended after the distractor. Conversely, for a distractor less salient than the target (Exp. 2), although behavioral interference was present, no attentional capture was observed. Together, these findings reveal two types of spatial-attentional dynamics in the auditory modality (distraction with and without attentional capture).

**Significance Statement:** Oftentimes, we find it hard to ignore the location of a salient sound that distracts us from our current tasks. Although, a common everyday experience, little is known about how spatial distraction unfolds at the neural level in the auditory modality. Using electrophysiological markers of attention allocations, we report comprehensive evidence of spatial attentional capture by a salient auditory distractor, indicating that attention is first misallocated to the distractor and only afterwards reallocated towards the target. Similar patterns were observed earlier only in vision and their discovery in the auditory modality indicates towards the existence of domain-general spatial attentional dynamics consistent across sensory modalities. Finally, we demonstrate that only a distractor more salient than the target reliably captures attention.

## Tracking the misallocation and reallocation of spatial attention towards auditory stimuli

People are constantly bombarded with a wide range of sensory experiences. To select and attend the important ones amongst multiple simultaneous stimuli is a massive computational challenge (Tsotsos, 1990; Bronkhorst, 2000). Likely as a side effect of these highly efficient selection mechanisms (Liesefeld et al., 2021), attention is sometimes misallocated towards a salient-but-irrelevant stimulus – leading to behavioral costs. Although almost exclusively studied in vision, this “problem” of *attentional capture* against the observers will appears also relevant for audition. Consider a typical office scenario from an auditory perspective– the sounds from mouse clicks or keyboard typing by a busy co-worker, grinding of the coffee machine, footsteps of someone walking by, or even the fan noise of an overheated computer. All these sounds are perceived simultaneously and emerge from various locations – although people actively pay attention to only a small subset of them. A knock on the office door (a salient event which is relevant to the office worker), would immediately draw attention to the door. However, if a bird pecked on the window or a co-worker accidentally dropped their coffee-mug (an irrelevant-but-salient event) it would also capture attention.

There is little research on such spatial attentional capture by auditory stimuli, but similar scenarios have been extensively studied with visual stimuli using the *additional-singleton paradigm* – where a salient-but-irrelevant stimulus (distractor) occurring simultaneously with a to-be-found target stimulus involuntarily captures attention under certain conditions (Hickey et al., 2006; Kiss et al., 2012; Burra and Kerzel, 2013; Gaspar et al., 2016; Liesefeld et al., 2022). To study the attentional dynamics involved in visual attentional capture, the N2pc component of the event-related potential (ERP), has proven highly useful. The N2pc component is a transient negative increase in activity over posterior electrode sites contralateral to the attended stimulus – and it is often used as a marker of spatial attention allocations (Eimer, 1996, 2014; Constant et al., 2023). It has been used to measure the timing of attention allocations (Töllner et al., 2011; Grubert and Eimer, 2016) and, in particular, to uncover the temporal dynamics of attentional capture by salient visual distractors (Liesefeld et al., 2017, 2022; Liesefeld and Müller, 2019). Strictly speaking, the notion of attentional capture implies a series of events, involving the initial misallocation of attention to the salient distractor, followed by re-allocating attention towards the target. The notion of sequential attention allocations also implies that the presence of distractors delays attention allocation to the target compared to when the distractor is absent. These spatio-temporal dynamics predict a very specific N2pc pattern: (a) the occurrence of a distractor N2pc – reflecting attentional capture by the distractor, (b) a delay in target N2pc when the distractor is present compared to distractor absence, and (c) a delay in target N2pc compared to the distractor N2pc (Liesefeld and Müller, 2019).

There are evidence of an auditory analog to the N2pc which has been termed *N2ac* (Gamble and Luck, 2011). The N2ac component is also a transient negative potential contralateral to the location of an attended stimulus but it is elicited by auditory stimuli and observed at more anterior electrode sites. This component is thought to be functionally similar to the N2pc component (e.g., Gamble and Woldorff, 2015; Lewald and Getzmann, 2015; Klatt et al., 2018). Thus, if it is possible to induce auditory attentional capture, we should be able to conceptually replicate the above-described pattern of spatio-temporal dynamics that has been taken as indicative of visual attentional capture. This would allow to further validate the functional interpretation of the N2ac component and to prove the existence of spatial attentional capture in the auditory domain. Most importantly, demonstrating such a relatively complex pattern of spatio-temporal attentional dynamics for the auditory modality in light of previous research in vision, would be strong evidence for more fundamental, modality-overarching principles of spatial dynamics of attentional (mis)allocation.

## Experiment 1

### Methods

In order to build a basic auditory search scene that would allow to disentangle various attentional dynamics, we adapted the design of Hickey et al. (2009). This design provides several advantages for our purposes. Hickey et al. used a visual search display with two objects (a square and a line), one stimulus being defined as the target while the other served as the distractor. Participants were asked to report on a feature of the target (e.g., whether the square target was a square or diamond – when the square is rotated 90°, or if the line target was a small or big line). Such tasks in which the target is defined by one feature (e.g., its shape) and participants are to report one of its features (e.g., its size) are called discrimination, categorization, or compound search tasks (Liesefeld et al., 2023) and are commonly employed in the additional-singleton paradigm (Theeuwes, 1991). Notably, by placing one stimulus on the midline and the other lateralized, Hickey et al. were able to disentangle attentional dynamics related to the respective lateralized stimulus, namely either the target or the distractor, in the presence of the midline stimulus. Just as the N2pc studied in Hickey et al., the N2ac employed here is a component emerging contralateral to the attended stimulus and stimuli on the midline consequently cannot elicit such lateralized activity. The additional advantage of the Hickey et al. (2009) design for our study is that only very few stimuli were presented simultaneously (see also Eimer, 1996; Hilimire et al., 2012). Presenting many identical (non-target) stimuli simultaneously (as would be the default for visual-search studies) is not feasible for studies using spatial sounds, because the auditory system cannot spatially segregate more than a few concurrently presented sounds, in particular if these stimuli are quite abstract and similar (Bregman, 1990), like those employed in typical visual-search displays.

On this background, the auditory scene employed in the present study consisted of just two clearly distinguishable sounds (sine waves vs. square-tone bursts) – where the instructions defined one as the target and the other as the distractor. Participants were required to report on each trial whether the target stimulus was of high frequency or low frequency indicated through pressing either of two keys.

#### Participants

Sixteen participants took part in Experiment 1 (median age: 26 years; range: 23 - 38 years, 9 males). This sample size is in the upper range of previous N2pc and N2ac studies and sufficient to detect effects of size *d*_z_ = 0.75 and above, with a probability of *1 −* β = .80 (α = .05, two-tailed). In both the experiments, participants reported normal hearing. Written informed consent was obtained from all the participants, and they received course credit or were paid for their participation. No participant had to be excluded. Procedures were approved by the ethics committee of the Department of Psychology and Pedagogics at Ludwig-Maximilians-Universität Munich.

#### Experimental Design

Stimulus presentation and response collection was controlled using PsychoPy (Peirce et al., 2019). The auditory search stimuli were presented using 3 studio sound monitors (Genelec 8020D, Genelec, Iisalmi, Finland) that were placed in the frontal field (see Figure 1a). One sound monitor was placed directly in front of the participants at 0° angle while the other two sound monitors were at approximately 40° to the left and right, respectively, at 110 cms from the participants. Participants were asked to rest their head on a chinrest throughout the experiment such that there are minimal head movements and distance of head from the sound source is kept constant.

**Figure 1.**
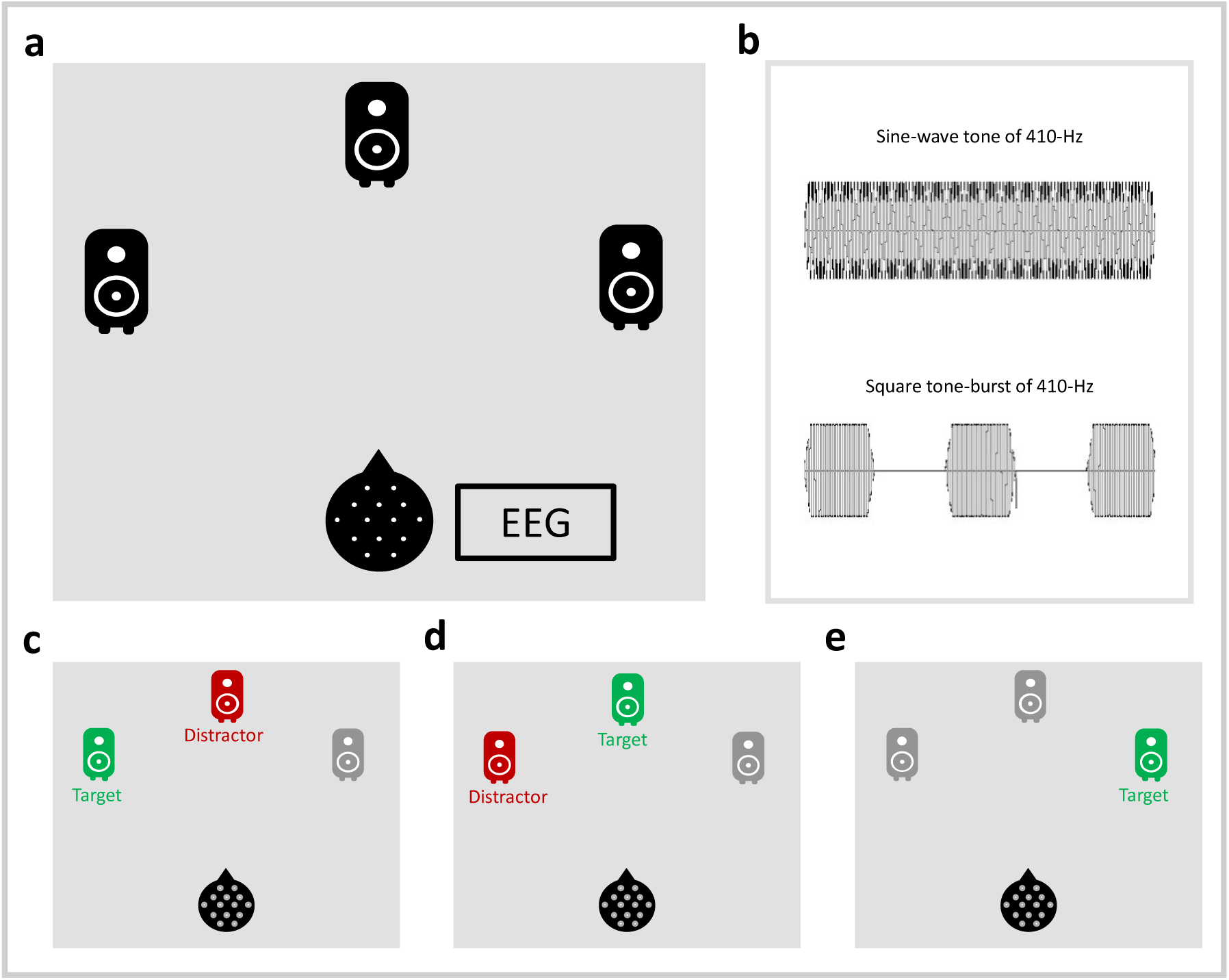
Auditory search scene. **(a)** Setup for the auditory search. **(b)** Representation of the waveform of the two kinds of sounds used. **(c)** Distractor-midline/target-lateral condition. **(d)** Distractor-lateral/target-midline condition. **(e)** Distractor-absent/target-lateral condition.

On each trial, either one (distractor-absent condition; target-only) or two (distractor-present condition) sounds were presented from different locations. The target sounds were sine-wave tones of either 440-Hz or 470-Hz. Participants’ task was to indicate whether the target was the high-frequency (i.e., the 470-Hz) or the low-frequency (the 440-Hz) tone in each trial. In distractor-present trials, an additional sound, from a different location than the target was presented. The distractor sounds were square tone-bursts (comprising of three 50-ms bursts separated by 50-ms of silence; see Figure 1b for a visual representation of the stimuli’s waveforms), which could either be of low-frequency (410-Hz) or of high-frequency (500-Hz). The temporal gaps in the distractor sounds render them more salient than the continuous sine-wave targets (Kayser et al., 2005). Both the distractor and the target sounds were presented for 250- ms at ∼70 dB(A) SPL, measured at the eardrum, using the miniDSP EARS device (miniDSP, Hong Kong, China). Participants were instructed to maintain eye fixation at a fixation cross presented on a monitor in front of them while performing the task. The monitor was placed such that it was below the sound monitor at 0° and did not obstruct the sound from it. Participants had to respond within 3000 ms after the onset of the sound (*response deadline*). In case of incorrect or delayed responses, the fixation cross changed to red or blue for 1000 ms, respectively. No feedback was provided for correct responses. The inter-trial interval was jittered between 800 ms and 1200 ms.

The main part of the experiment consisted of 20 blocks of 108 trials each (2160 trials in total), with 720 distractor-absent trials and 1440 distractor-present trials. An additional practice block was provided at the beginning of the experiment, which consisted of 108 unanalyzed trials.

The experiments were designed to isolate attentional processing of target and distractor through lateralized event-related potentials (ERPs) – specifically the N2ac component (Gamble and Luck, 2011). The design follows a similar logic as used to dissect the attentional dynamics of target and distractor processing in visual search using the N2pc component (e.g., Liesefeld et al., 2022). The *distractor-midline/target-lateral* condition (Figure 1c) contained a lateralized target with a distractor on the midline and served to isolate target-related activity. The *distractor-lateral/target-midline* condition (Figure 1d) contained a lateralized distractor with a target on the midline and served to isolate distractor-related activity. The *distractor-absent/target-lateralized* condition (Figure 1e) contained only a lateralized target without a distractor and served as a baseline for target related activity in the absence of a distractor. To avoid any bias in target or distractor location, we also ran a condition with the distractor on one and the target on the other side, which confounds target- and distractor-related activity and is therefore not further discussed here. Within each block, the three display configurations as well as frequency of distractor and target were completely balanced and randomized in distractor-present trials, which were randomly intermixed with distractor-absent trials in which target frequency was balanced and randomized as well.

#### Electrophysiological recording and analysis

The EEG was recorded via 60 pre-amplified Ag/AgCl electrode positioned according to the international 10-10 system. Horizontal ocular artifacts were monitored via two additional electrodes on the outer canthi of both eyes. All impedances were kept below 15 kΩ. Signals were amplified (250-Hz low-pass filter, 10-s time constant; BrainAmp DC, BrainProducts) and sampled at 1000 Hz. EEG data were processed with custom-written Matlab scripts using functions from EEGLAB (Delorme and Makeig, 2004), ERPLAB (Lopez-Calderon and Luck, 2014) and the ‘latency.m’ function from (Liesefeld, 2018).

Signals were re-referenced offline to the average of both mastoids. A 0.5-Hz high-pass and a 40-Hz low-pass FIR filter (EEGLAB default) were applied, after which independent-component analysis (ICA) was ran on the signal. The ICA components representing blinks or horizontal eye-movements were then removed from the continuous EEG data. After this, the data were segmented into epochs from -200 to 800 ms relative to the search-stimuli onset and baseline-corrected relative to the pre-stimulus interval. The trials with artifacts in the analyzed channels (FC5/6; voltage steps larger than 50 μV per sampling point, activity changes <0.5 μV within a 500-ms time window, or absolute amplitude exceeding ±80 μV; equal to *M* = 1.81%, *min* = 0.15% and *max* = 7.63% of trials in Exp. 1 and *M* = 5.47%, *min* = 0.06% and *max* = 18.33% of trials in Exp. 2), or incorrect responses were excluded (at least 355 trials remaining per individual in each condition after trial exclusion). The electrode positions FC5/6 were chosen (instead of the anterior electrode cluster used in Gamble & Luck, 2011) as N2ac has been found to be most prominent at these locations in previous research (Lewald and Getzmann, 2015; Lewald et al., 2016).

In order to determine the analysis window for the component of interest, the on- and offsets of the strongest component of the respective polarity was identified as the time points where the ERP in the grand-average difference wave (GA) crossed 30% of the component’s peak amplitude. The 50%-area latency was used for the component latency estimation, where the component area was bounded by the ERP, a threshold set at 30% of the respective component’s peak amplitude searched in a common time window encompassing GA on- and offsets of all analyzed components (Liesefeld et al., 2017, 2022; Liesefeld, 2018). Amplitudes were defined as the mean activity in a 30-ms window centered on each component’s 50%-area latency.

#### Statistical analyses

For amplitudes, we report *P*_perm_ values obtained from a permutation method modelled after Sawaki et al. (2012). The procedure followed is the same as that used in (Liesefeld et al., 2022). For each of the 10,000 permutations, *n*_L_ and *n*_R_ trials were randomly assigned from the respective display-configuration to the left and right ERP. Here, *n*_L_ and *n*_R_ represents the number of trials that went into the respective original individual ERPs. The GA waveform was built from these, and the signed amplitude was extracted in the time-range of 0 – 500 ms. The *P*_perm_ values indicate the proportion of runs in the random permutation that yielded an amplitude larger than or equal to that of the correct assignment of left and right trials. In other words, *P*_perm_ can be interpreted as the probability of observing a value larger or equal to the observed amplitude merely due to random fluctuations.

Pairwise comparisons between distractor-absent and distractor-present conditions for median correct reaction times (RTs) and error rates were performed using Wilcoxon signed-rank test (two-tailed comparisons). Results graphs show the mean of medians and within-participant confidence intervals (Cousineau and O’Brien, 2014). Additionally, we report Bayes factors (*BF*s), for the Bayesian Wilcoxon signed-rank test (result based on data augmentation algorithm with 5 chains of 10,000 iterations), quantifying evidence for the alternative over the null hypothesis (*BF*₁₀). BFs were calculated using the standard JZS Cauchy prior with a scale factor of *r* = √2/2. To classify the strengths of evidence through the Bayes Factors, we used Jeffreys’s criterion (van Doorn et al., 2021) – which states that for the alternative hypothesis, *BF*s between 1 and 3 are weak evidence, BFs between 3 and 10 are moderate evidence, and BFs greater than 10 are strong evidence. Following these criteria, 1/3 (= 0.33) < *BF*_10_ < 1 is considered weak evidence for the null hypothesis, 1/10 (= 0.10) < *BF*_10_ < 1/3 (= 0.33) is considered as moderate evidence for the null hypothesis, and *BF*_10_ < 1/10 (= 0.10) is considered strong evidence for the null hypothesis. All analyses were performed using JASP v0.17.2 (JASP Team, 2023) and using custom scripts in Matlab. The data, analysis scripts and results are available on OSF (https://osf.io/qr86k/?view_only=e50053edad0044039903867f6228ca24).

### Results

#### Distractor interference

Compared to distractor-absent trials (*M* = 531 ms, *Mdn* ± *MAD* = 517 ± 79 ms), distractor presence (*M* = 596 ms, *Mdn* ± *MAD* = 577 ± 93 ms) significantly delayed responses by 65 ms, *W* = 136.00, *p* < .001, *r* = 1.00, *BF*_10_ = 36919.71. The *r* = 1.00 means that each individual participant was delayed by distractor presence. The error rates for the distractor-present trials (*M* = 9.3 %, *Mdn* ± *MAD* = 8.55 ± 4.45 %), were also significantly higher (by 3.17%) from the distractor-absent trials (*M* = 6.13 %, *Mdn* ± *MAD* = 5.94 ± 3.92 %), *W* = 121.00, *p* =.004, *r* = .78, *BF*_10_ = 32.61. Thus, overall, we observe a significant distractor interference effect in both RTs and error rates (see Figure 2a).

**Figure 2.**
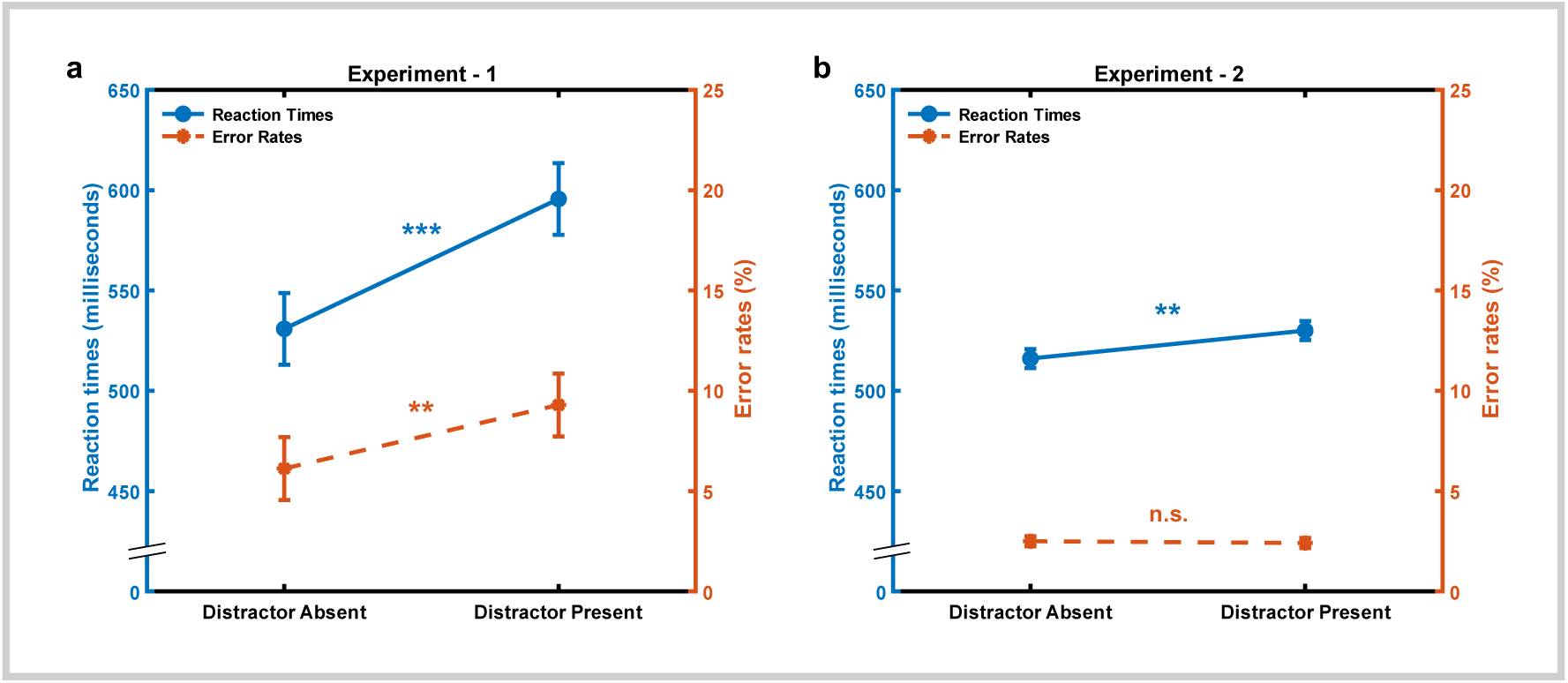
Distractor interference in Experiments 1 and 2. **(a)** Mean of the median RTs and error rates for the distractor absent and distractor present conditions in Experiment 1. **(b)** RTs and error rates in Experiment 2. Error bars represent 95% within-participant confidence intervals. n.s. p > .05, **p < .01, ***p < .001.

#### Distractor presence delays attention allocation towards the target

As expected, spatial attention was (eventually) directed to the location of the target irrespective of whether the distractor was present or absent. This was evidenced by the N2ac component which was observed for the target in both the distractor absent (*P*_perm_ < .001) and the distractor midline (*P*_perm_ < .001) condition (see Figure 3a). Importantly, distractor presence (*M* = 271.06 ms; *Mdn* ± *MAD* = 267.5 ± 13.5 ms) delayed the target N2ac by 56.87 ms (*W* = 124.5, *p* = .004, *r* = .831, *BF*_10_ = 58.90), compared to distractor absence (*M* = 214.19 ms; *Mdn* ± *MAD* = 225 ± 33 ms). This pattern of result indicates that the salient distractor indeed delayed the allocation of attention to the target. Such delay in attention allocation to the target due to a distractor is well documented in visual search using the N2pc component (e.g., Liesefeld et al., 2017, 2022). This delay is comparable to that induced by visual distractors in Liesefeld et al. (2017); 59 ms and Liesefeld et al. (2022); 70 – 88 ms.

**Figure 3.**
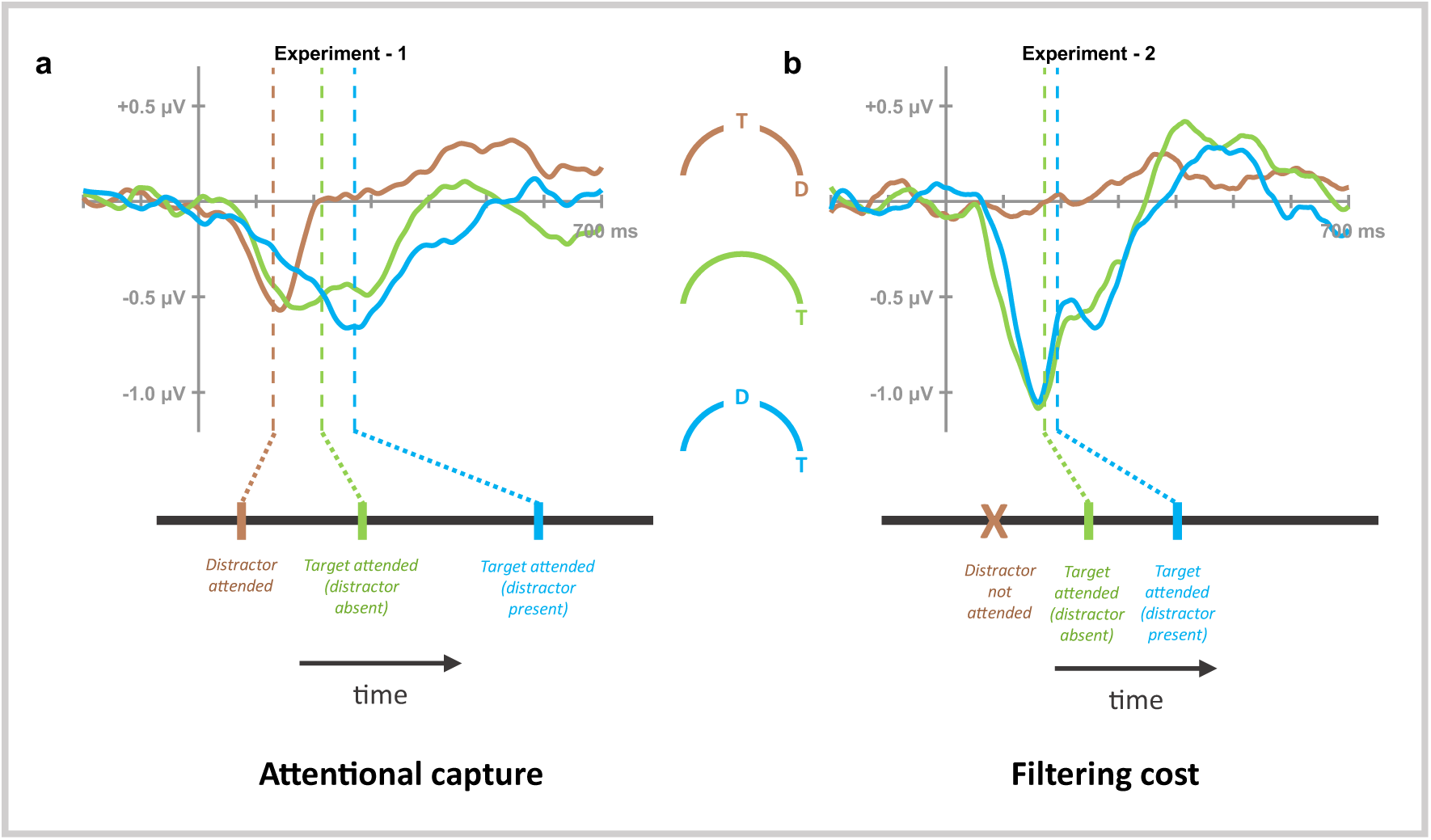
Tracking the dynamics of attentional capture and filtering cost using N2ac. Bottom panels shows the expected temporal pattern of the N2ac during attentional capture and filtering cost, while the top panel demonstrates the exact pattern of results as obtained from experiment 1 **(a)** and 2 **(b)** using the N2ac component. Vertical dashed-lines in the ERPs (top panels) indicate mean 50%-area latencies. A smoothing filter (Savitzky–Golay filter; order = 3, frame length = 101) was applied to this and subsequent waveforms before visualization to improve the visibility of effects.

#### Attentional capture by a salient distractor: Distractor N2ac

The interference due to the distractor (as observed through the behavioral results and the delay in target N2ac due to distractor presence) could be due to the distractor entering the competition for attention and either winning this competition – thereby capturing attention, or (barely) failing to win the competition – thus only delaying the attention towards the target (“non-spatial filtering costs”; Becker, 2007; Liesefeld et al., 2019). The expected temporal dynamics corresponding to the two mentioned attentional mechanisms as informed through visual search is illustrated in the bottom panel of Figure 3. If attention is indeed captured by the salient distractor, then we should observe a significant N2ac towards the lateral distractor. To resolve for this, we analyzed the distractor-lateral / target-midline condition and observed a significant N2ac (*P*_perm_ = .026) for the distractor (Figure 3a), thus indicating that the distractor indeed captured attention.

#### Attention shifts from distractor to the target

As a final criterion to consider the observed interference as attentional capture, we predicted that spatial attention as indicated by the N2ac first goes to the distractor and only afterwards moves on to the target (see Liesefeld and Müller, 2019). To examine this, we compared the distractor N2ac latency in the distractor-lateral/target-midline condition to the latency of the N2ac in the distractor-midline/target-lateral condition. Indeed, the N2ac towards the distractor (*M* = 129.62 ms; *Mdn* ± *MAD* = 136 ± 15 ms) was 141.44 ms earlier than the N2ac towards the target (*M* = 271.06 ms; *Mdn* ± *MAD* = 267.5 ± 13.5 ms), *W* = 136.00, *p* < .001, *r* = 1.00, *BF*_10_ = 6811.39 (Figure 3a). This indicates that attention shifts from the distractor to the target. The time required to shift attention from one sound to the other is within the estimates for the time required for relocating attention in visual search (100–150 ms; Woodman and Luck, 2003) and is even a bit longer than that empirically observed for visual targets and distractors by Liesefeld et al. (2017); 100 ms and Liesefeld et al. (2022); 99 – 119 ms.

## Experiment 2

### Methods

One possible critique of the findings in Experiment 1 could be that, although completely in line with attentional capture, the N2ac pattern observed is due to low-level sensory imbalances caused by the presentation of a stimulus in one auditory hemifield and not due to attentional processes. To rule out this possibility and to also examine distractor interference without attentional capture (filtering costs), we performed a control experiment where we swap the roles of target and the distractor from Exp. 1 (Gaspar and McDonald, 2014; Barras and Kerzel, 2017; Gaspelin et al., 2023). If the N2ac pattern observed in Exp. 1 was due to sensory imbalances in the auditory hemifield, we would expect to observe the exact same pattern in Exp. 2. If, however, the N2ac in Exp. 1 really indicates attentional capture, we should not replicate it when switching target and distractor identities. This is because the square-tone bursts employed as a distractor in Exp. 1 and as a target in Exp. 2 are much more salient than the sine-wave sounds, such that it should be much easier to select the target *and* to ignore the distractor (see, Gaspar & McDonald, 2014; Zehetleitner et al., 2013) in this experiment.

A new sample of 16 participants (median age: 24 years; range: 19 - 44 years; 8 males) took part in Experiment 2. The general setup and the task design were similar to that of Experiment 1, except that the target- and distractor-defining features were swapped (while the reported features, i.e., the base frequencies remained same). That is, the target was now a square tone-burst (three 50-ms tone burst separated by 50-ms silence in between) of either 440-Hz (low-frequency) or 470-Hz (high-frequency). The distractors were continuous sine tone of either 410-Hz (low-frequency) or 500-Hz (high-frequency). Both target and distractors were presented for 250 ms as in Exp. 1. Here again, participants were instructed to report whether the target was of high-frequency or low-frequency. Importantly, the switch of target- and distractor-defining features was supposed to render the target more salient than the distractor.

### Results

#### Behavioral costs due to distractors

Responses on distractor-present trials (*M* = 530 ms, *Mdn* ± *MAD* = 501 ± 62 ms) were significantly delayed (by 14 ms; see Figure 2b) compared to distractor-absent trials (*M* = 516 ms, *Mdn* ± *MAD* = 497 ± 67 ms), *W* = 130.00, *p* < .001, *r* = .91, *BF*_10_ = 354.56. Error rates showed no significant difference (see Figure 2b) between distractor-present (*M* = 2.42 %, *Mdn* ± *MAD* = 1.32 ± 0.76 %) and distractor-absent trials (*M* = 2.5 %, *Mdn* ± *MAD* = 1.74 ± 1.04 %), *W* = 38.00, *p* = .222, *r* = -.37, *BF_10_* = 0.48.

#### Delay in attention allocation to target due to distractor presence

As in Experiment 1, here also attention was allocated to the target location – as indicated by an N2ac component that occurred when the distractor was present (*P*_perm_ < .001) or absent (*P*_perm_ < .001). There was a delay of 22.25 ms (*W* = 105.50, *p* = .011, *r* = .76, *BF*_10_ = 15.76) for the target N2ac (see Figure 3b) when the distractor was present (*M* = 193.81; *Mdn* ± *MAD* = 185.5 ± 27.5) compared to when the distractor was absent (*M* = 171.56; *Mdn* ± *MAD* = 164 ± 10.5), however, this delay was much smaller than that observed for Exp. 1.

#### No evidence for attentional capture by the distractor: Absence of distractor N2ac

Unlike in Experiment 1, where the distractor clearly captured attention as indicated by a distractor N2ac, in Experiment 2 we do not observe a statistically reliable N2ac to the distractor (*P*_perm_ = .570; see Figure 3b). In the absence of electrophysiological evidence for attentional capture by the less salient distractor the distractor-presence effect on behavior and target N2ac latency is best interpreted as non-spatial filtering costs (in contrast to attentional capture in Experiment 1; see Introduction).

#### N2ac latency as measure of relative salience

As discussed earlier, the N2ac component is believed to be the auditory analog of the lateralized N2pc component (sometimes also referred to as PCN; Töllner et al., 2008). Stimulus salience is known to modulate the latency of the N2pc component elicited by a visual-search target, such that with increasing salience N2pc latency decreases (Töllner et al., 2011). Comparing distractor-absent target N2ac latencies across the two experiments can therefore serve to confirm that the square-tone burst is more salient than the sine tone (when presented in isolation). As expected, in the distractor-absent/target-lateralized condition we indeed see a significant decrease in the target N2ac latency (see Figure 4) by 42.63 ms for Experiment 2 (more salient target) compared to that of Experiment 1 (less salient target; *U* = 197.00, *p* = .010, *r* = .54, *BF_10_* = 2.25)^*^, thereby, confirming that the sine-tone stimulus is indeed less salient than the square-tone-burst stimulus.

**Figure 4.**
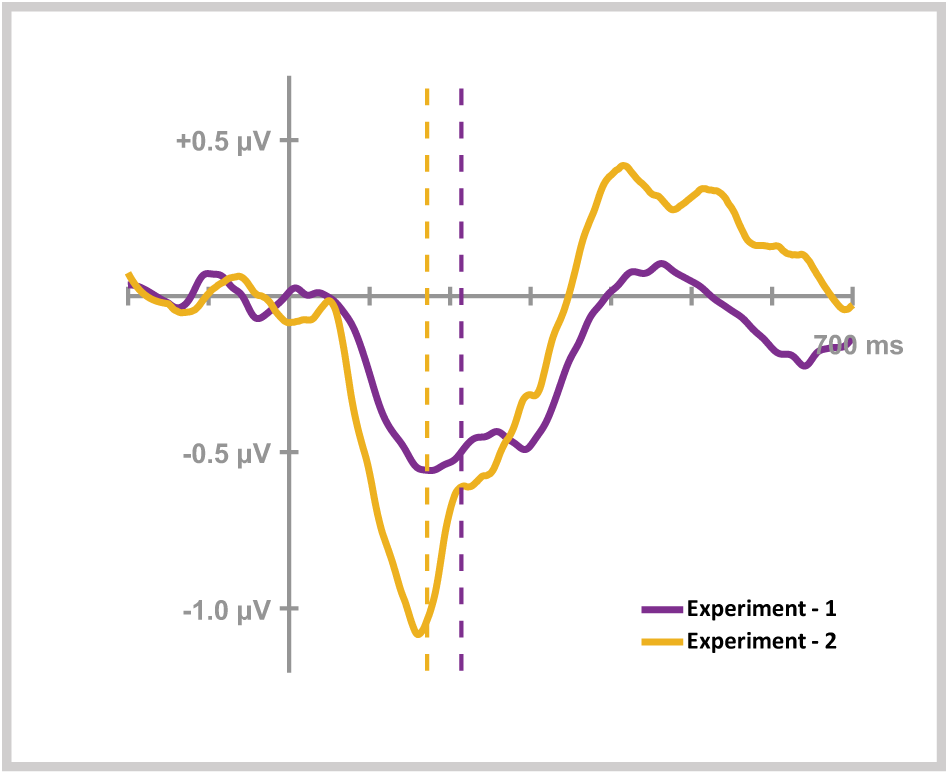
N2ac latency indicates relative salience of stimuli. The target N2ac latency when distractor is absent decreases for Exp. 2 compared to Exp. 1, indicating an increased relative salience of the target stimuli in Exp. 2 (square-tone bursts) compared to Exp. 1 (sine tones).

## Discussion

The present study investigated the dynamics of spatial attention in auditory search using the N2ac component. An auditory distractor of higher salience than the target (Exp. 1) captured spatial attention as indicated by an N2ac to the distractor. After this initial misallocation, attention was reallocated to the target as indicated by an ensuing target N2ac, which was delayed compared to a distractor-absent condition. This signature N2ac pattern is the most comprehensive ERP evidence for attentional capture, as explained by (Liesefeld and Müller, 2019). In contrast, an auditory distractor less salient than the target in Experiment 2 failed to capture attention, although it still delayed attention allocation towards the target, a phenomenon that is described as non-spatial filtering costs in visual search. This is the first report of both these spatial attentional phenomena in the auditory domain.

### Indication of a domain-general attention allocation mechanism

The N2pc component has been extensively studied in visual search over the last decades and has been established as a marker for the spatial allocation of attention in vision (Eimer, 1996, 2014; Constant et al., 2023). Similar lateralized ERP components have been discovered for other modalities, such as the N2ac component (Gamble and Luck, 2011) for audition (which we study here) and the N2cc component (also known as the CCN; Katus & Eimer, 2019; Tsai et al., 2023) for tactile modality. Both components are believed to be functional analogs of the N2pc component in their respective sensory modalities – used to index spatial allocation of attention.

The temporal dynamics of mis-allocation and consequent reallocation of spatial attention due to salient distractors has been earlier demonstrated only in vision through the N2pc component (Liesefeld et al., 2017, 2022). That the N2ac component indicates similar attentional dynamics for audition, not only strengthens the functional interpretation of the N2ac component as an auditory analog to the N2pc component used to measure spatial attentional allocations in the auditory modality, but also points towards the existence of domain-general spatio-temporal attentional dynamics – indicating a more fundamental principle of attentional misallocation and reallocation across all sensory modalities (see also Shinn-Cunningham, 2008 for a related discussion).

### Differences in spatial attentional allocation for detection and discrimination tasks

In the original N2ac study (Gamble and Luck, 2011), the target N2ac emerged only when the target was accompanied by a distractor, whereas it emerged with and without a concurrently presented distractor in the present study (see Figure 3). This discrepancy is likely due to differences in the task designs employed in the two studies. Gamble & Luck (2011) employed a detection task, wherein participants were tasked with determining the presence or absence of the target in each trial. In contrast, we employed a discrimination task, which involves the differentiation of a specific target feature (reported feature). It is well established that detection tasks impose comparatively lower attentional demands compared to discrimination tasks (Lavie, 2005), and that detection might occur in a pre-attentive manner (Treisman and Gelade, 1980; Luck et al., 1997). In scenarios involving target detection, an exhaustive focal-attentional engagement with the target stimuli might not be obligatory, particularly when the saliency of the target is high (which is indeed the case for the targets used in Gamble & Luck, 2011). Instead, target presence can in principle be deduced from an inhomogeneity on the priority map induced by a salience signal emitted from the target (Müller et al., 2004; Liesefeld et al., 2016). This signal can inform the correct response without necessitating any spatial attentional allocation and thus without eliciting an N2ac. Only when there are competing salience signals in search-detection tasks focal-attentional analysis might be required for confirming target presence (Hoffman, 1979). In contrast, discrimination of the target’s reported feature (high vs. low frequency in our case) in search-discrimination tasks might make a spatial shift of attention towards the target mandatory, yielding a pronounced N2ac component even in the absence of any competing stimulus.

### Absence of a distractor suppression (P_D_) component

Along with the target N2pc component, Hickey et al. (2009) observed a P_D_ component – reflecting inhibition of the salient visual distractor (see Gaspelin et al., 2023 for a comprehensive review). Interestingly, we do not observe any reliable P_D_ in the current study. Given the similarity in our design with Hickey et al. (2009) – except for the modality difference, one could expect to observe signatures of spatial suppression for distractor-lateral/target-midline condition. Such a P_D_ component – indicative of spatial suppression has been recently reported for auditory distractors (Lunn et al., 2023). The study reported a transient positive deflection in anterior electrodes contralateral to the auditory distractor in the time range of 100 – 300 ms – comparable to a P_D_ commonly observed for visual distractors. In their study, attention was not captured, so that their P_D_ would be interpreted as proactive or stimulus-triggered suppression (see Liesefeld et al., 2023) that potentially serves to avoid attentional capture (Gaspelin and Luck, 2018, 2019). We did not observe any indication of a positivity in this time range in our Experiment 2. When attention is captured – as apparently was the case in our Exp. 1 – such a P_D_ should only occur *after* the distractor N2ac, indicating reactive suppression (see Gaspelin et al., 2023; Liesefeld et al., 2023); comparable to what has been observed in visual search (Liesefeld et al., 2017, 2022; Liesefeld and Müller, 2019). Although we indeed do observe a small positive deflection contralateral to the distractor occurring in a reasonable time-range of 400 – 600 ms in Experiment 1 (Figure 3a), this component did not reach statistical significance. Future research might examine whether and under which circumstances reliable pre- and post-distractor-N2ac P_D_s (indicative of distractor suppression before or after attentional capture) emerge in auditory search.

### Differences between audition and vision

As already pointed out before Gamble and Luck (2011), the lesser degree of contralaterality of the auditory system compared to the visual system might explain why the N2ac (here and in other studies) is smaller in peak amplitude (0.5 – 1 µV) than the typical N2pc (1 - 2 µV). Another reason for small peak amplitudes might be that the N2acs observed so far are less tightly locked to the sound onset and therefore smeared out across time (therefore of lower peak amplitude and longer in duration than the typical N2pc). In fact, stretched-out N2pcs of smaller peak amplitude and higher latency (i.e., smeared out N2pcs) have been observed when they likely reflect the second attention allocation (Liesefeld et al., 2017) or when the target is harder to find (Töllner et al., 2011; Dowdall et al., 2012). In the present study, the N2acs in Experiment 1 (where the target was relatively harder to find) are also relatively less in peak amplitude and longer in duration (i.e., more smeared) than those observed in Experiment 2 (where the target was relatively easier to find). Thus, pointing towards similar factors (in their respective modalities) influencing the amplitude and duration of both the N2ac and N2pc components.

Also note that the N2ac seems to emerge earlier than the typical N2pc. In the ERP graphs shown above (Figure 3), for example, the N2ac seems to emerge (onset) already at around 70 ms, whereas the typical N2pc does not occur before around 130 ms (see Liesefeld et al., 2017, 2022, for examples from the same lab, and Hickey et al., 2009, for search displays of comparable complexity). In line with this observation, simple reaction times to auditory stimuli are typically 20-60 ms shorter than to visual stimuli (Lewald and Guski, 2003; Jose and Gideon Praveen, 2010) which is likely the result of faster physiological processing time in the auditory system than the visual system (Stein and Meredith, 1993). In fact, already the very first steps of perceptual processing (the transduction process) are of a magnitude faster in audition compared to vision (Recanzone, 2009). Thus, it is not surprising that also attentional dynamics occur earlier when triggered by auditory events compared to visual events. This would be the case even if the attentional process itself is highly comparable or even functionally identical.

## Acknowledgements

This project was funded by the German Research Association (DFG) via the RTG 2175 “Perception in Context and its Neural Basis”.

## Footnotes

* Results from Frequen st and Bayesian Mann Whitney U test.

## Notes

**Conflict of interest statement**: We have no conflict of interest to disclose.

### Competing Interest Statement

The authors have declared no competing interest.

## References

Barras C, Kerzel D (2017) Salient-but-irrelevant stimuli cause attentional capture in difficult, but attentional suppression in easy visual search. Psychophysiology 54:1826–1838.

Becker SI (2007) Irrelevant singletons in pop-out search: Attentional capture or filtering costs? J Exp Psychol Hum Percept Perform 33:764–787.

Bregman AS (1990) Auditory Scene Analysis: The Perceptual Organization of Sound. The MIT Press. Available at: https://direct.mit.edu/books/book/3887/Auditory-Scene-AnalysisThe-Perceptual-Organization [Accessed July 19, 2023].

Bronkhorst A (2000) The Cocktail Party Phenomenon: A Review of Research on Speech Intelligibility in Multiple-Talker Conditions. Acta Acust United Acust 86:117–128.

Burra N, Kerzel D (2013) Attentional capture during visual search is attenuated by target predictability: Evidence from the N2pc, Pd, and topographic segmentation. Psychophysiology 50:422–430.

Constant M, Mandal A, Asanowicz D, Yamaguchi M, Gillmeister H, Kerzel D, Luque D, Pesciarelli F, Fehr T, Mushtaq F, Pavlov YG, Liesefeld HR (2023) A multilab investigation into the N2pc as an indicator of attentional selectivity: Direct replication of Eimer (1996). Available at: https://psyarxiv.com/3472y/ [Accessed May 13, 2023].

Cousineau, O’Brien F (2014) Error bars in within-subject designs: a comment on Baguley (2012). Behav Res Methods 46:1149–1151.

Delorme A, Makeig S (2004) EEGLAB: an open source toolbox for analysis of single-trial EEG dynamics including independent component analysis. J Neurosci Methods 134:9–21.

Dowdall JR, Luczak A, Tata MS (2012) Temporal variability of the N2pc during efficient and inefficient visual search. Neuropsychologia 50:2442–2453.

Eimer M (1996) The N2pc component as an indicator of attentional selectivity. Electroencephalogr Clin Neurophysiol 99:225–234.

Eimer M (2014) The neural basis of attentional control in visual search. Trends Cogn Sci 18:526– 535.

Gamble ML, Luck SJ (2011) N2ac: An ERP component associated with the focusing of attention within an auditory scene. Psychophysiology 48:1057–1068.

Gamble ML, Woldorff MG (2015) The temporal cascade of neural processes underlying target detection and attentional processing during auditory search. Cereb Cortex 25:2456– 2465.

Gaspar JM, Christie GJ, Prime J, Jolicœur P, Mc onald JJ (2016) Inability to suppress salient distractors predicts low visual working memory capacity. Proc Natl Acad Sci 113:3693– 3698.

Gaspar JM, McDonald JJ (2014) Suppression of Salient Objects Prevents Distraction in Visual Search. J Neurosci 34:5658–5666.

Gaspelin N, Lamy D, Egeth HE, Liesefeld HR, Kerzel D, Mandal A, Müller MM, Schall JD, Schubö A, Slagter HA, Stilwell BT, van Moorselaar D (2023) The Distractor Positivity Component and the Inhibition of Distracting Stimuli. J Cogn Neurosci:1–23.

Gaspelin N, Luck SJ (2018) Combined Electrophysiological and Behavioral Evidence for the Suppression of Salient Distractors. J Cogn Neurosci 30:1265–1280.

Gaspelin N, Luck SJ (2019) Inhibition as a Potential Resolution to the Attentional Capture Debate. Curr Opin Psychol 29:12–18.

Grubert A, Eimer M (2016) All set, indeed! N2pc components reveal simultaneous attentional control settings for multiple target colors. J Exp Psychol Hum Percept Perform 42:1215– 1230.

Hickey C, Di Lollo V, McDonald JJ (2009) Electrophysiological Indices of Target and Distractor Processing in Visual Search. J Cogn Neurosci 21:760–775.

Hickey C, McDonald JJ, Theeuwes J (2006) Electrophysiological Evidence of the Capture of Visual Attention. J Cogn Neurosci 18:604–613.

Hilimire MR, Hickey C, Corballis PM (2012) Target resolution in visual search involves the direct suppression of distractors: Evidence from electrophysiology: Target resolution and distractor suppression. Psychophysiology 49:504–509.

Hoffman JE (1979) A two-stage model of visual search. Percept Psychophys 25:319–327.

Jose S, Gideon Praveen K (2010) Comparison between Auditory and Visual Simple Reaction Times. Neurosci Med 2010 Available at: http://www.scirp.org/journal/PaperInformation.aspx?PaperID=2689 [Accessed August 18, 2023].

Katus T, Eimer M (2019) The N2cc component as an electrophysiological marker of space-based and feature-based attentional target selection processes in touch. Psychophysiology 56:e13391.

Kayser C, Petkov CI, Lippert M, Logothetis NK (2005) Mechanisms for Allocating Auditory Attention: An Auditory Saliency Map. Curr Biol 15:1943–1947.

Kiss M, Grubert A, Petersen A, Eimer M (2012) Attentional Capture by Salient Distractors during Visual Search Is Determined by Temporal Task Demands. J Cogn Neurosci 24:749–759.

Klatt L-I, Getzmann S, Wascher E, Schneider D (2018) The contribution of selective spatial attention to sound detection and sound localization: Evidence from event-related potentials and lateralized alpha oscillations. Biol Psychol 138:133–145.

Lavie N (2005) Distracted and confused?: Selective attention under load. Trends Cogn Sci 9:75– 82.

Lewald J, Getzmann S (2015) Electrophysiological correlates of cocktail-party listening. Behav Brain Res 292:157–166.

Lewald J, Guski R (2003) Cross-modal perceptual integration of spatially and temporally disparate auditory and visual stimuli. Cogn Brain Res 16:468–478.

Lewald J, Hanenberg C, Getzmann S (2016) Brain correlates of the orientation of auditory spatial attention onto speaker location in a “cocktail-party” situation. Psychophysiology 53:1484–1495.

Liesefeld HR (2018) Estimating the Timing of Cognitive Operations With MEG/EEG Latency Measures: A Primer, a Brief Tutorial, and an Implementation of Various Methods. Front Neurosci 12:765.

Liesefeld HR et al. (2023) Terms of debate: Consensus definitions to guide the scientific discourse on visual distraction. Available at: https://psyarxiv.com/4b2gk/ [Accessed June 9, 2023].

Liesefeld HR, Liesefeld AM, Müller HJ (2019) Distractor-interference reduction is dimensionally constrained. Vis Cogn 27:247–259.

Liesefeld HR, Liesefeld AM, Müller HJ (2021) Attentional capture: An ameliorable side-effect of searching for salient targets. Vis Cogn 29:600–603.

Liesefeld HR, Liesefeld AM, Müller HJ (2022) Preparatory Control Against Distraction Is Not Feature-Based. Cereb Cortex 32:2398–2411.

Liesefeld HR, Liesefeld AM, Töllner T, Müller HJ (2017) Attentional capture in visual search: Capture and post-capture dynamics revealed by EEG. NeuroImage 156:166–173.

Liesefeld HR, Moran R, Usher M, Müller HJ, Zehetleitner M (2016) Search efficiency as a function of target saliency: The transition from inefficient to efficient search and beyond. J Exp Psychol Hum Percept Perform 42:821–836.

Liesefeld HR, Müller HJ (2019) Distractor handling via dimension weighting. Curr Opin Psychol 29:160–167.

Lopez-Calderon J, Luck SJ (2014) ERPLAB: an open-source toolbox for the analysis of event-related potentials. Front Hum Neurosci 8:213.

Luck SJ, Girelli M, McDermott MT, Ford MA (1997) Bridging the gap between monkey neurophysiology and human perception: An ambiguity resolution theory of visual selective attention. Cognit Psychol 33:64–87.

Lunn J, Berggren N, Ward J, Forster S (2023) Irrelevant sights and sounds require spatial suppression: ERP evidence. Psychophysiology 60:e14181.

Müller H, Krummenacher J, Heller (2004) imension specific intertrial facilitation in visual search for pop out targets: Evidence for a top down modulable visual short term memory effect. Vis Cogn 11:577–602.

Peirce J, Gray JR, Simpson S, MacAskill M, Höchenberger R, Sogo H, Kastman E, Lindeløv JK (2019) PsychoPy2: Experiments in behavior made easy. Behav Res Methods 51:195–203.

Recanzone GH (2009) Interactions of auditory and visual stimuli in space and time. Hear Res 258:89–99.

Sawaki R, Geng JJ, Luck SJ (2012) A Common Neural Mechanism for Preventing and Terminating the Allocation of Attention. J Neurosci 32:10725–10736.

Shinn-Cunningham BG (2008) Object-based auditory and visual attention. Trends Cogn Sci 12:182–186.

Stein BE, Meredith MA (1993) The Merging of the Senses. MIT Press.

Theeuwes J (1991) Cross-dimensional perceptual selectivity. Percept Psychophys 50:184–193.

Töllner T, Gramann K, Müller HJ, Kiss M, Eimer M (2008) Electrophysiological markers of visual dimension changes and response changes. J Exp Psychol Hum Percept Perform 34:531– 542.

Töllner T, Zehetleitner M, Gramann K, Müller HJ (2011) Stimulus Saliency Modulates Pre-Attentive Processing Speed in Human Visual Cortex. PLOS ONE 6:e16276.

Treisman AM, Gelade G (1980) A feature-integration theory of attention. Cognit Psychol 12:97– 136.

Tsai S-Y, Nasemann J, Qiu N, Töllner T, Müller HJ, Shi Z (2023) Little engagement of attention by salient distractors defined in a different dimension or modality to the visual search target. Psychophysiology:e14375.

Tsotsos JK (1990) Analyzing vision at the complexity level. Behav Brain Sci 13:423–445.

van Doorn J et al. (2021) The JASP guidelines for conducting and reporting a Bayesian analysis. Psychon Bull Rev 28:813–826.

Woodman GF, Luck SJ (2003) Serial deployment of attention during visual search. J Exp Psychol Hum Percept Perform 29:121–138.

Zehetleitner M, Koch AI, Goschy H, Müller HJ (2013) Salience-Based Selection: Attentional Capture by Distractors Less Salient Than the Target. PLOS ONE 8:e52595.

